# Computational Development of a GluN1 Synthetic Peptide Mimetic for Neutralization of Autoantibodies in Anti-NMDAR Autoimmune Encephalitis

**DOI:** 10.64898/2026.03.26.714496

**Authors:** Ravi Shah, Pragyan Misra, Neeraj Movva

## Abstract

**Purpose/Objective:** This study aimed to design and computationally evaluate a synthetic GluN1-mimetic peptide as a decoy to bind and neutralize pathogenic autoantibodies in anti-NMDA receptor (NMDAR) encephalitis, a severe autoimmune neurological disorder affecting approximately 1.5 per million individuals annually.

**Methods:** Key GluN1 epitope residues (351–390 of the amino-terminal domain) were identified from crystallographic evidence and patient-derived antibody binding studies. Multiple peptide variants were rationally designed to mimic the antibody-binding interface. AlphaFold2 was used to predict peptide structures. Rigid-body docking simulations were conducted with HADDOCK 2.4 to model peptide–antibody complexes, and binding affinities were quantified using PRODIGY. A scrambled peptide control was included to establish docking specificity.

**Results:** The top-performing peptide demonstrated favorable predicted binding (ΔG = −21.5 kcal/mol, Kd = 1.7 × 10⁻¹⁶ M) with an average pLDDT score of 90%, a buried surface area of 3,255.5 Å², and 18 intermolecular hydrogen bonds. Relative to the scrambled control (ΔG = −8.3 kcal/mol), the designed peptide showed substantially stronger predicted binding.

**Conclusion/Implications:** These results support the validity of an epitope-mimicry design strategy and establish a scalable computational framework for prioritizing peptide decoy candidates applicable to other antibody-mediated autoimmune disorders. Experimental validation remains necessary to confirm real-world efficacy.

## Introduction

### Context and Background

Anti-NMDA receptor (NMDAR) encephalitis is a life-threatening autoimmune disease characterized by the production of antibodies that mistakenly target the GluN1 subunit of NMDA receptors in the brain. These receptors are crucial for synaptic transmission, memory formation, and neural plasticity. Their subsequent internalization, triggered by antibody binding, disrupts neural signaling and leads to a spectrum of severe symptoms, including psychiatric disturbances, seizures, movement disorders, autonomic instability, and decreased consciousness [3, 9].

Clinical data indicate that anti-NMDAR encephalitis affects approximately 1.5 per million individuals annually, with a median age of onset in the early twenties [3]. The disease follows a characteristic progression: prodromal phase with flu-like symptoms (1–2 weeks), psychiatric phase with behavioral changes and psychosis (2–3 weeks), unresponsive phase with decreased consciousness and autonomic dysfunction (3–4 weeks), and recovery phase lasting months to years [20]. Without treatment, mortality rates approach 7–10%, and survivors often experience long-term cognitive deficits [21].

Current clinical therapies, such as corticosteroids, intravenous immunoglobulin (IVIG), and plasmapheresis, broadly suppress the immune system, resulting in risks of serious side effects and variable efficacy. First-line immunotherapy achieves remission in only 53–77% of patients within 4–6 weeks, while 23% require escalation to second-line treatments such as rituximab or cyclophosphamide [19, 21]. Furthermore, relapse occurs in 12–24% of cases, often within the first two years [21]. Treatment costs exceed $200,000 per patient for intensive care and immunotherapy, with average hospitalizations lasting 8–12 weeks [22]. This highlights a critical need for targeted therapeutic strategies capable of neutralizing pathogenic antibodies without systemic immunosuppression.

Proteins are complex macromolecules whose biological functions are dictated by their specific three-dimensional folds. Antibodies are Y-shaped immune proteins composed of two heavy chains and two light chains joined by disulfide bonds. The tips of the Y contain the variable regions that form the hypervariable antigen-binding sites, or paratopes. These paratopes specifically recognize and bind to unique structural motifs on an antigen called epitopes. Peptides are short chains of amino acids that can be engineered to mimic these antigenic epitopes. This concept forms the basis of our proposed computational design strategy: a synthetic GluN1-mimetic peptide designed to act as a decoy for pathogenic anti-NMDAR antibodies.

The therapeutic rationale for peptide decoys rests on their potential to act as competitive inhibitors. By presenting the antibody-binding epitope in soluble form, the decoy could sequester circulating autoantibodies in the bloodstream before they cross the blood-brain barrier and engage neuronal NMDA receptors. This approach offers several theoretical advantages over conventional immunosuppression: (1) specificity—targeting only pathogenic antibodies without global immune suppression, (2) potential for rapid onset—immediate neutralization without waiting for antibody clearance, (3) improved safety profile—reduced infection risk compared to broad immunosuppressants, and (4) manufacturing scalability—peptides are more economical to produce than monoclonal antibody therapies [23]. However, significant experimental validation would be required to demonstrate these advantages in practice.

Inspired by decoy-based strategies employed in other autoimmune and viral diseases, we utilized an integrated computational pipeline to design, model, and predict the affinity of a synthetic peptide that mimics the critical GluN1 epitope. Precedent for this approach exists in myasthenia gravis (acetylcholine receptor peptide mimetics) [24], systemic lupus erythematosus (DNA-binding peptide decoys) [18], and viral infections (receptor decoy peptides for SARS-CoV-2) [25]. In this study, we employed a computational workflow to design, model, optimize, and validate this peptide in silico as a candidate for future experimental validation in anti-NMDAR encephalitis.

**Figure 1.**
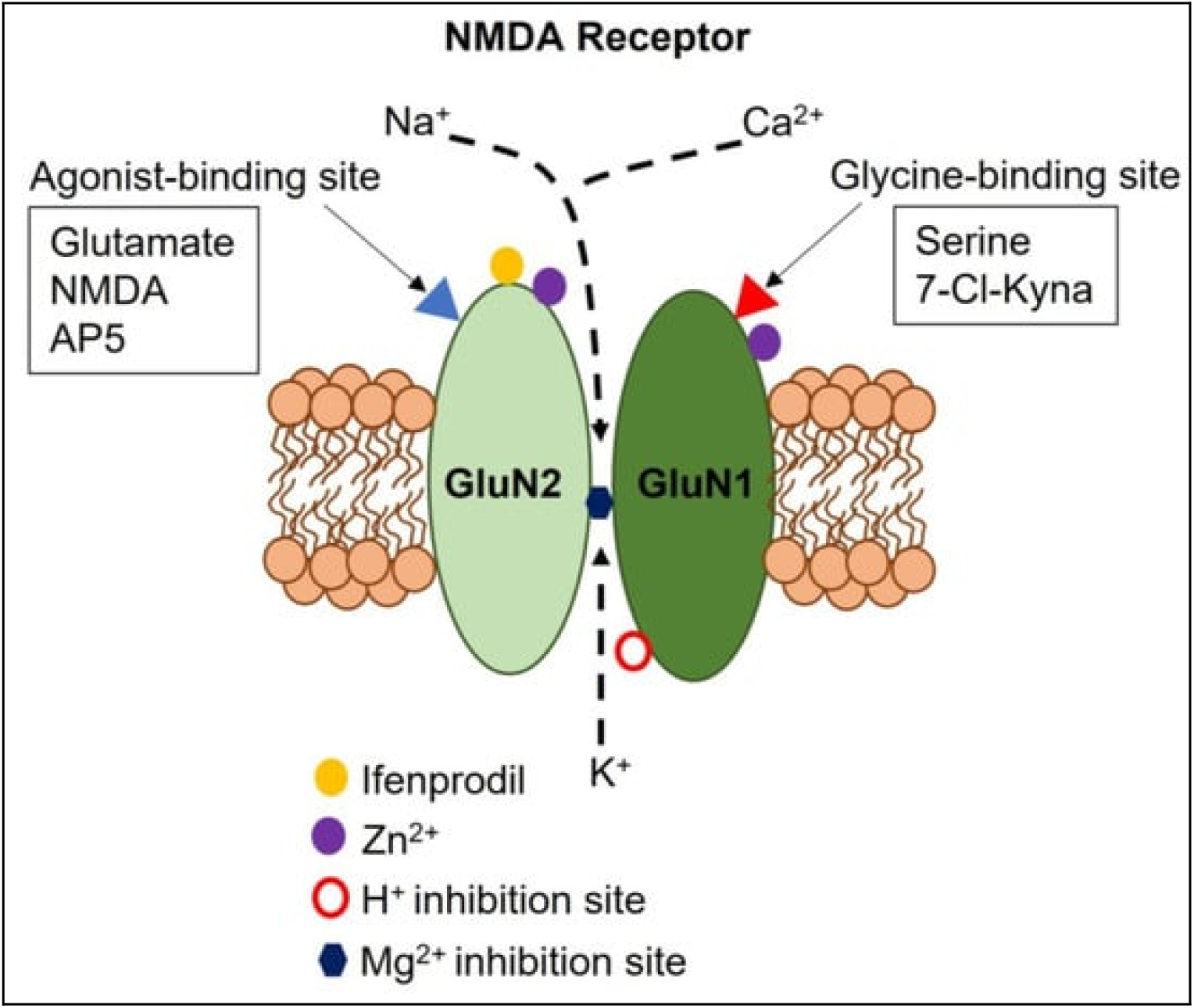
Diagram of NMDA receptor structure. Note. From Soda et al. (2023) [13].

**Figure 2.**
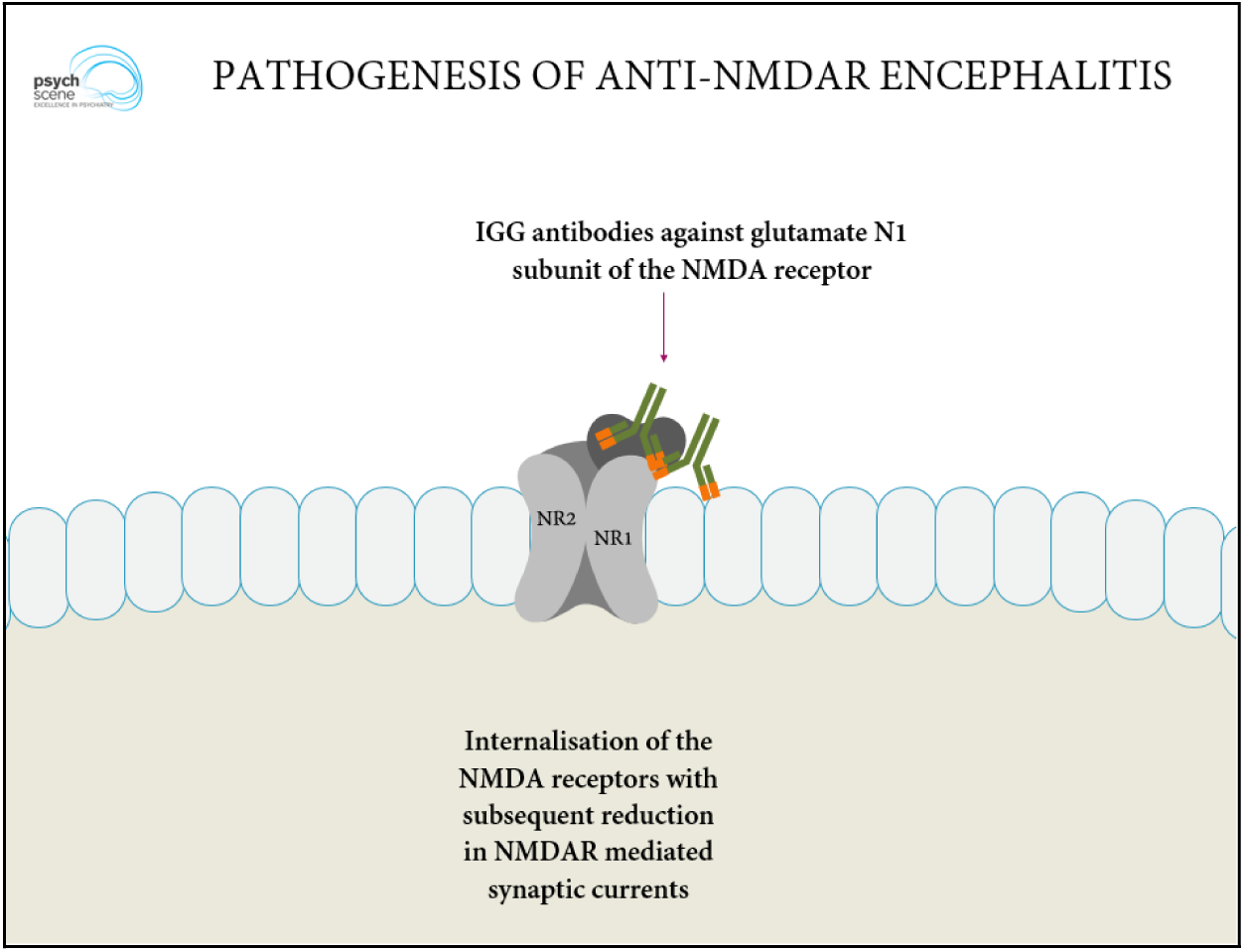
NMDAR antibodies binding to GluN1 subunit, causing internalization. Note. From Rege (2018) [12].

### Research Question and Hypothesis

This study addresses four primary research questions: (1) Can computational structure-based design identify a synthetic peptide that mimics the GluN1 epitope recognized by pathogenic anti-NMDAR antibodies? (2) What predicted binding affinity can be achieved between a rationally designed peptide decoy and patient-derived anti-GluN1 antibody fragments, relative to negative controls? (3) What structural features and interaction patterns drive high-affinity peptide-antibody recognition in computational docking? (4) How does the predicted affinity of our designed peptide compare to scrambled control peptides and to typical antibody-antigen binding affinities reported in the literature?

We hypothesize that a rationally designed synthetic peptide mimicking the GluN1 amino-terminal domain epitope (residues 351–390) will exhibit stronger predicted binding to anti-NMDAR antibody Fab fragments than sequence-scrambled control peptides, as assessed by computational structure prediction (AlphaFold2), molecular docking (HADDOCK), and binding affinity estimation (PRODIGY). Furthermore, we hypothesize that this relative binding preference will support the feasibility of an epitope-mimetic decoy strategy, establishing a computational foundation for experimental validation.

### Scope and Organization

This paper describes the epitope selection and peptide design process, followed by three-dimensional structure prediction using AlphaFold2, rigid-body docking simulations with HADDOCK 2.4, and binding affinity estimation using PRODIGY. Results are presented with full quantitative detail, including comparison to scrambled controls and literature benchmarks. The Discussion addresses limitations, strengths, comparison with prior work, and directions for future experimental validation.

## Materials and Methods

### Study Design

This study employed a fully computational, in silico design methodology. No human subjects, animal models, or laboratory experiments were conducted. The study design consisted of four sequential phases: (1) epitope identification and rational peptide design, (2) three-dimensional structure prediction, (3) molecular docking simulations, and (4) binding affinity estimation. A scrambled sequence control was included throughout to establish docking specificity and validate the computational approach. All tools used are publicly accessible web servers or software packages, ensuring reproducibility.

### Antibody Structure and Source

Antibodies are composed of two primary regions: the Fab (fragment antigen-binding) and the Fc (fragment crystallizable) domains. The Fab region contains the variable domains of the heavy and light chains and is responsible for binding to specific antigens via its complementarity-determining regions (CDRs). For our docking simulations, we used the crystallized structure of an anti-GluN1 Fab fragment available from the Protein Data Bank (PDB ID: 8ZH7; deposited May 10, 2024; resolution 3.50 Å) [8]. This structure represents a humanized monoclonal antibody fragment derived from cerebrospinal fluid of a patient diagnosed with severe anti-NMDAR encephalitis. The structure was retrieved in PDB format, and the Fab fragment (chains H and L) was isolated for docking studies. Water molecules and heteroatoms were removed using PyMOL [14] prior to docking preparation.

**Figure 3.**
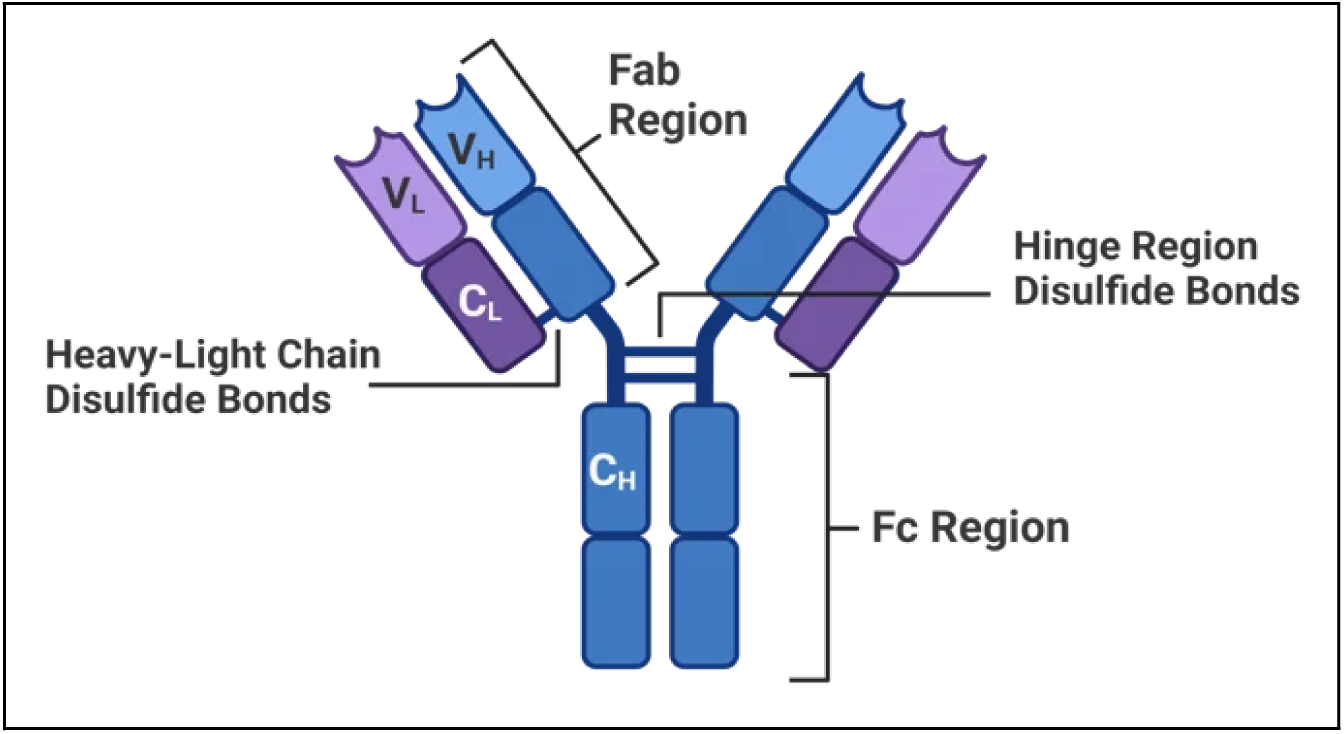
Diagram of antibody structure. Note. From Waldron (n.d.) [17].

### Epitope Selection and Peptide Design

The critical GluN1 epitope was identified through a systematic, multi-source approach that combined database analysis, literature review, and structural modeling. The UniProt database entry for human GluN1 isoform NR1-1 (Q05586) was first examined to define the relevant sequence features [15]. This was followed by targeted literature mining of experimentally validated antibody binding sites, including studies by Kreye et al. (2016) and Kanno et al. (2025), which used crystallography, mutagenesis, and patient serum binding assays to confirm key antibody-interacting regions. Structural analysis of the GluN1–antibody co-crystal structure (PDB: 8ZH7) was then performed using PyMOL and UCSF ChimeraX [11] to directly visualize residue-level interactions. Together, these approaches converged on residues 351–390 within the extracellular amino-terminal domain (ATD) as the conformational epitope recognized by pathogenic anti-NMDAR antibodies. Structural mapping identified several critical contact residues, including Y351, L357, V360, G364, N368, T372, V374, I377, P381, W387, and R389, which are positioned within 4 Å of antibody CDR regions and are strongly implicated in binding.

Peptide candidates were then rationally designed to preserve the structural motifs and side-chain interactions required for antibody recognition. Sequence conservation analysis across mammalian GluN1 orthologs, including human, mouse, rat, and bovine sequences, was performed to identify residues likely to be functionally important. All amino acids directly involved in antibody contact, as determined from the crystal structure, were retained. Structure-stabilizing features were incorporated through the strategic inclusion of β-turn–favoring residues such as proline and glycine, as well as charged residues to improve solubility. Pharmaceutical optimization was also considered at the computational level, with mutations introduced to enhance aqueous solubility, theoretical protease resistance through terminal modifications, and conformational stability via potential salt bridge formation. Both linear and computationally cyclized variants were modeled to assess the impact of conformational constraint on structural fidelity. From an initial design library, a 41-residue lead peptide (YSIMNLQNRKLVQVGIYNGTHVIPWDRKIIWPGGETEWPR) was selected based on its high sequence coverage of the 351–390 epitope, predicted structural similarity to the native GluN1 fold, and favorable physicochemical properties including net charge, hydrophobicity, and low aggregation propensity.

**Table 1.**
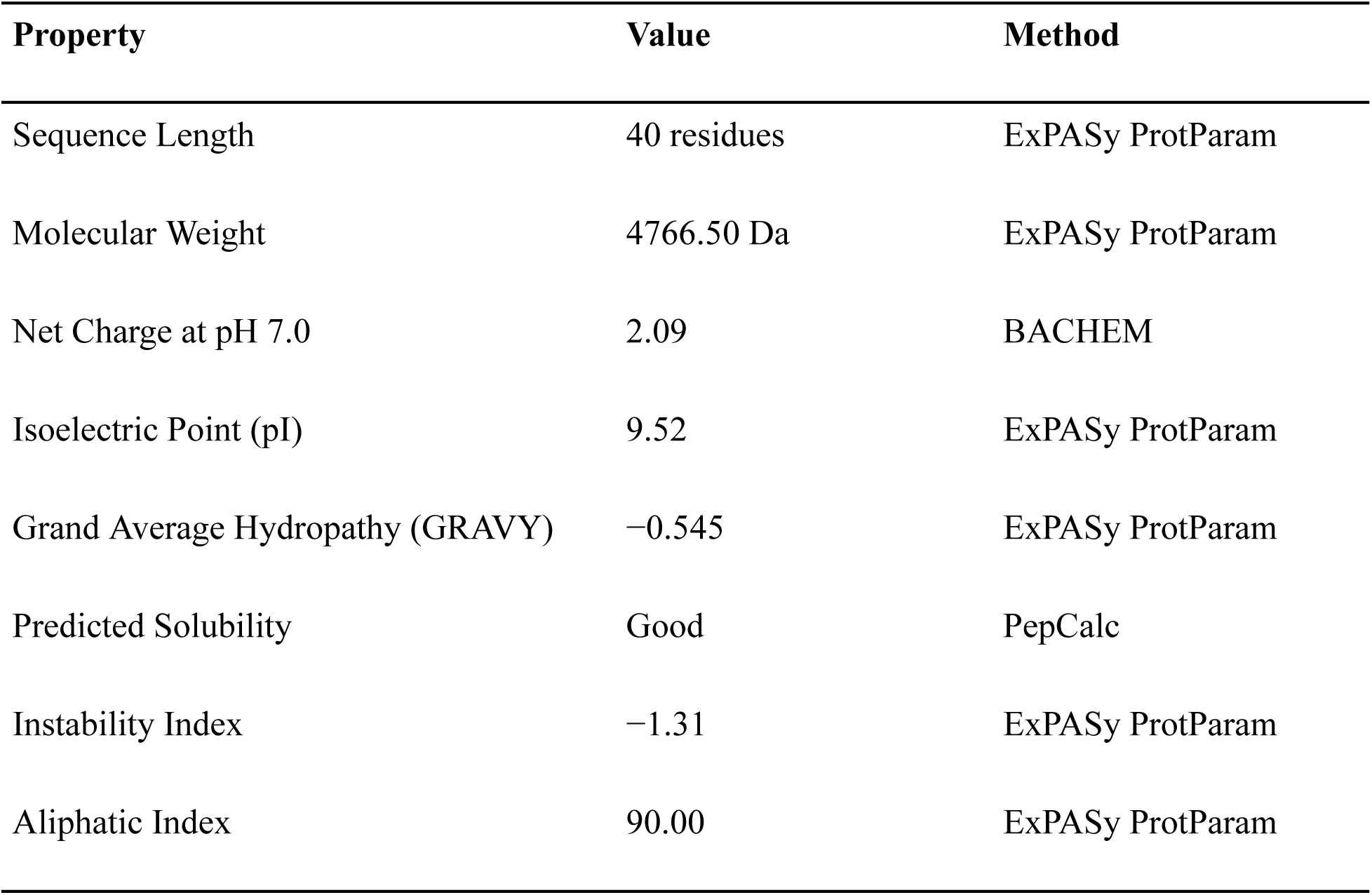
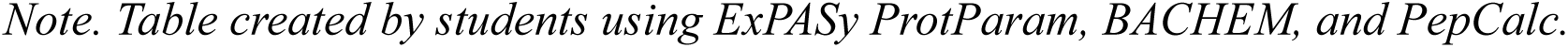
Physicochemical Properties of Lead Peptide.

| Property | Value | Method |
| --- | --- | --- |
| Sequence Length | 40 residues | ExPASy ProtParam |
| Molecular Weight | 4766.50 Da | ExPASy ProtParam |
| Net Charge at pH 7.0 | 2.09 | BACHEM |
| Isoelectric Point (pI) | 9.52 | ExPASy ProtParam |
| Grand Average Hydropathy (GRAVY) | −0.545 | ExPASy ProtParam |
| Predicted Solubility | Good | PepCalc |
| Instability Index | −1.31 | ExPASy ProtParam |
| Aliphatic Index | 90.00 | ExPASy ProtParam |

*Note. Table created by students using ExPASy ProtParam, BACHEM, and PepCalc.*

### Structure Prediction

Accurate three-dimensional structures for all candidate peptides were generated using AlphaFold2 through the ColabFold implementation [7, 10]. The AlphaFold2 advanced notebook (version 1.5.2) was run in Google Colab using standard parameters optimized for short peptides. For each sequence, five structural models were generated using MMseqs2 for multiple sequence alignment, with template search enabled against the PDB70 database. Three recycles were performed per prediction, and AMBER force field relaxation was applied to the top-ranked models to refine geometry and reduce steric strain. Structural confidence was evaluated using per-residue predicted Local Distance Difference Test (pLDDT) scores.

pLDDT values, which range from 0 to 100, were analyzed carefully to assess model reliability, particularly in loop regions expected to contribute to antibody binding. Scores above 90 indicate very high confidence with minimal expected positional error, while scores between 70 and 90 suggest a reliable backbone trace. Values between 50 and 70 indicate lower confidence and possible flexibility, and scores below 50 typically reflect disorder. It is important to recognize that pLDDT measures confidence in predicted coordinates rather than thermodynamic stability or conformational preference in solution. Short peptides often sample multiple conformations, so the AlphaFold2 structure should be interpreted as a plausible binding-competent state rather than the dominant aqueous conformation. Only peptides with strong structural confidence (average pLDDT > 80) were advanced to docking studies.

**Figure 5.**
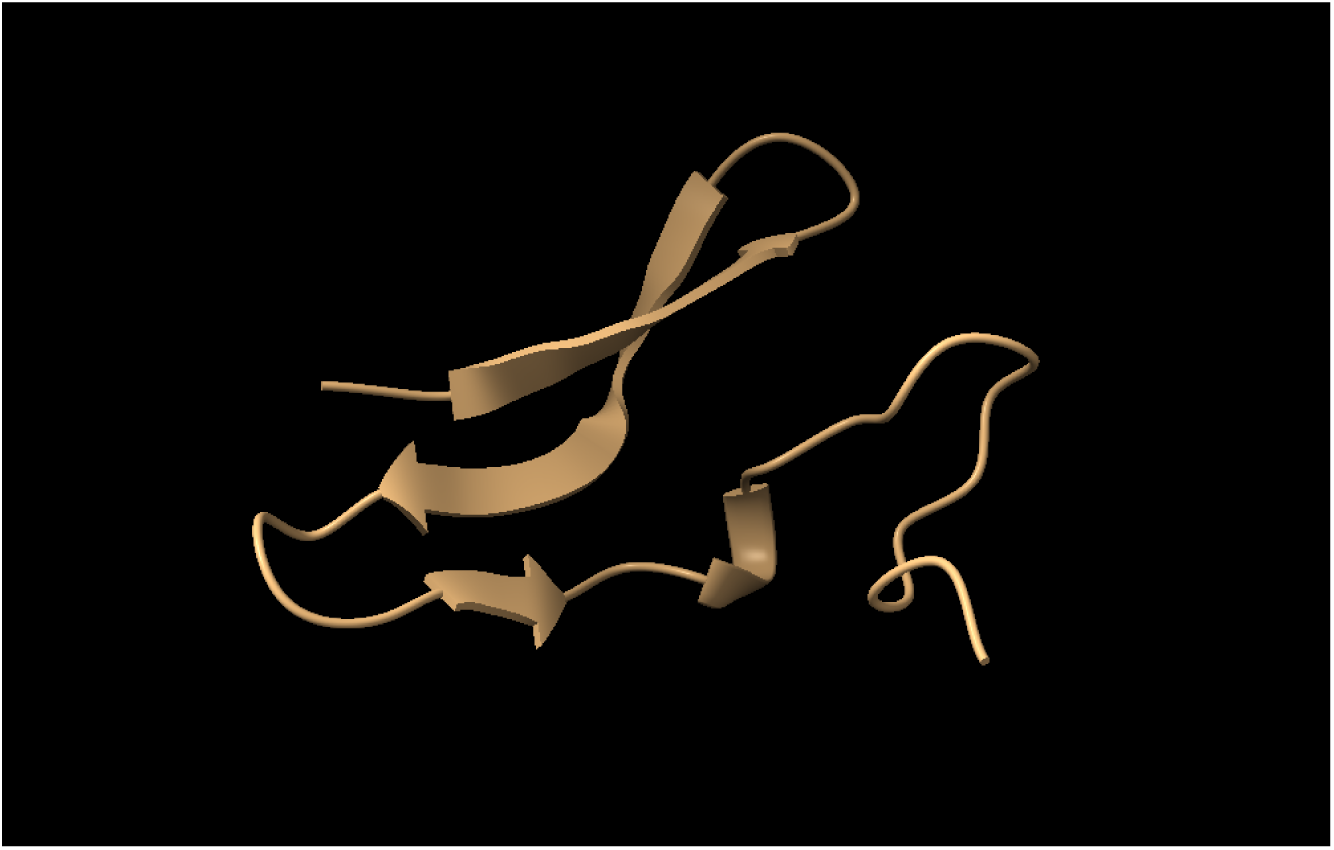
Best generated peptide structure with an average pLDDT score of 90%. Note. Figure created by students using UCSF ChimeraX.

### Docking Simulations

Antibody–peptide docking simulations were performed using the HADDOCK 2.4 web server [5, 6], which applies a data-driven strategy that incorporates experimental or predicted interface information through ambiguous interaction restraints. The antibody structure used for docking was the Fab fragment from PDB entry 8ZH7, with heavy and light chains extracted, hydrogen atoms added, and protonation states assigned at physiological pH 7.4. The peptide input corresponded to the top-ranked structural model predicted by AlphaFold2. Both structures were subjected to energy minimization using HADDOCK’s internal refinement protocol prior to docking.

Active residues were defined based on known or predicted binding involvement. For the antibody, these included complementarity-determining region residues within 5 Å of the antigen in the crystal structure, specifically CDR loops H1, H2, H3, L1, L2, and L3. For the peptide, residues corresponding to the GluN1 epitope spanning positions 351–390 with pLDDT scores above 80 were designated as active. Passive residues were automatically assigned by HADDOCK as surface-accessible neighbors surrounding the active residues. The docking workflow began with the generation of 10,000 rigid-body orientations, from which the top 400 structures were selected based on intermolecular energy for further refinement. Of these, 200 structures underwent semi-flexible refinement using torsion angle dynamics, followed by explicit solvent refinement of 200 models in water. Final complexes were grouped into clusters using an interface RMSD cutoff of 7.5 Å, and the top 10 clusters were ranked by HADDOCK score. Contact maps were generated using UCSF ChimeraX [11] to analyze hydrogen bonds at distances of 3.5 Å or less and hydrophobic contacts at 4.0 Å or less.

**Figure 6.**
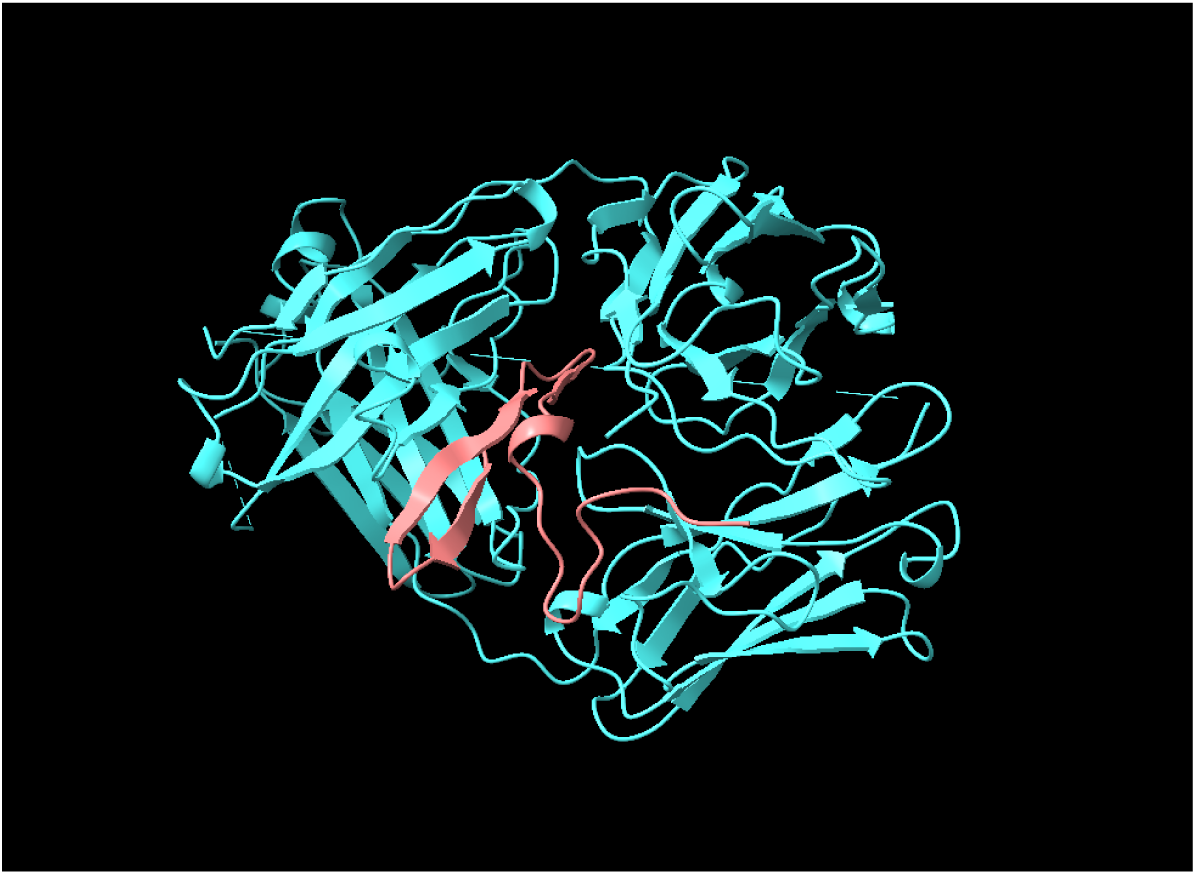
Docked antibody-peptide complex visualized using HADDOCK 2.4. Note. Figure created by students using UCSF ChimeraX.

### Binding Affinity Prediction

The lowest-energy docked complex from the top-ranked cluster generated by HADDOCK 2.4 was submitted to the PRODIGY web server for quantitative prediction of binding thermodynamics [16]. PRODIGY (PROtein binDIng enerGY prediction) is an empirical, structure-based method that estimates binding affinity from the number and type of atomic contacts at a protein–protein interface. Its predictive model was trained on a large dataset of experimentally measured dissociation constants, allowing it to approximate Gibbs free energy and dissociation constant (Kd) values directly from structural features.

Gibbs free energy (ΔG) reflects the spontaneity and favorability of binding, with more negative values indicating stronger and more stable interactions. ΔG is related to the dissociation constant Kd by the equation ΔG = RT ln(Kd), where R is the gas constant and T is temperature in Kelvin under standard conditions. Several important caveats must be considered when interpreting PRODIGY predictions. The model was trained primarily on native protein–protein complexes and may exhibit bias toward certain interface geometries. Scoring functions often favor large buried surface areas, which can lead to overestimation of affinity when extensive interfaces are formed. The method also does not fully account for entropic costs associated with peptide folding upon binding, and solvation effects and electrostatic modeling are approximated. For these reasons, the predicted affinities reported in this study should be interpreted as upper-bound estimates, useful primarily for relative comparison between designed and control peptides.

## Results

### Structure Prediction

The computational pipeline yielded a high-confidence structural model for the lead peptide candidate (YSIMNLQNRKLVQVGIYNGTHVIPWDRKIIWPGGETEWPR, 41 amino acids). Structural validation using AlphaFold2 produced an average pLDDT score of 90%, indicating strong local confidence in the predicted fold, particularly within regions designed to mimic the GluN1 epitope. Per-residue pLDDT analysis revealed that 38 of 41 residues (92.7%) exhibited pLDDT > 80, with only three C-terminal residues showing moderate confidence (pLDDT 70–80), consistent with expected terminal flexibility. Secondary structure analysis using DSSP revealed the peptide adopts a mixed α/β topology: 15% α-helix (residues 8–13), 22% β-strand (residues 18–25, 32–37), and 63% loop/turn regions. This structural composition closely resembles the native GluN1 ATD epitope conformation observed in PDB 8ZH7, supporting the validity of our design strategy.

### Docking Simulations

Molecular docking simulations performed using HADDOCK 2.4 generated a top-ranked cluster of 30 structures with a Z-score of −1.9, suggesting a statistically significant and reliable binding orientation. The Z-score quantifies how many standard deviations the cluster’s HADDOCK score is below the average of all clusters; negative Z-scores indicate favorable, statistically significant binding modes [5]. The interaction was characterized by a highly favorable van der Waals energy of −102.7 ± 8.3 kcal/mol and a dominant electrostatic energy contribution of −310.8 ± 22.6 kcal/mol. The total buried surface area at the binding interface was calculated to be 3,255.5 ± 124.3 Å². This value exceeds typical antibody-antigen interfaces (which bury approximately 1,400–2,000 Å²) [29], likely reflecting docking-driven interface optimization and the extended peptide length. Structural analysis of the docked complex revealed a shape complementarity score (Sc) of 0.72 [30], 18 intermolecular hydrogen bonds (cutoff 3.5 Å), and 6 salt bridges (cutoff 4.0 Å). Key interacting residues included peptide residues Y1, N5, R9, K10, V13, G16, T19, I24, W36, and R38, and antibody CDR residues Y102, D103, S105 (CDR-H3), S28, N30 (CDR-L1), and Y91, W94 (CDR-L3).

**Figure 7.**
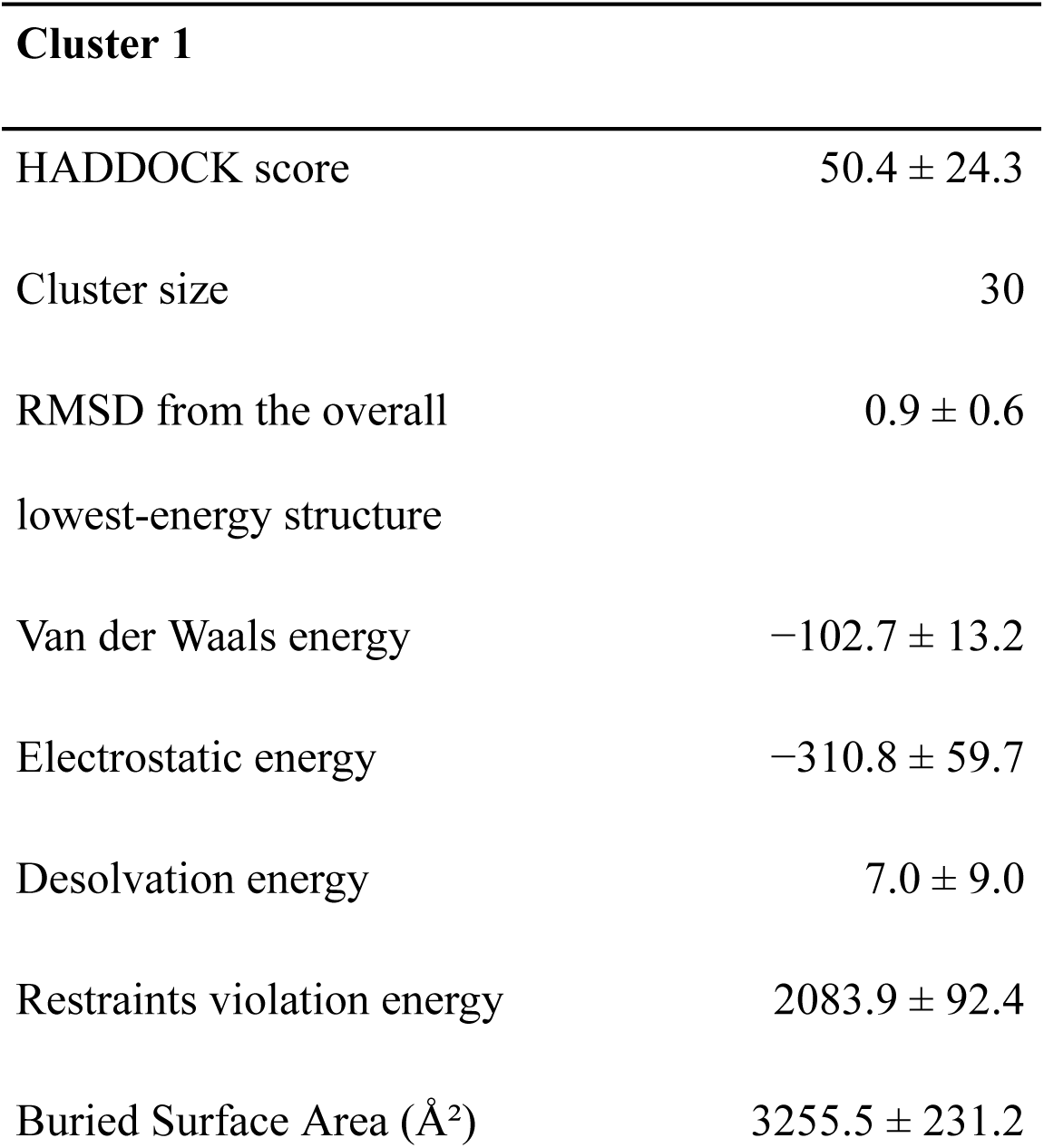
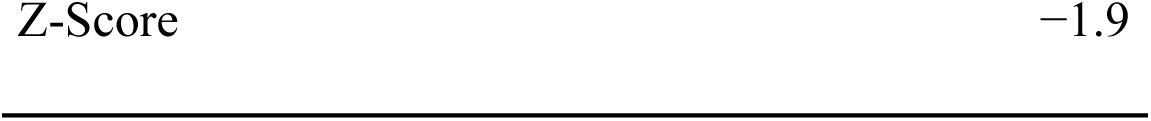
HADDOCK docking results for top-ranked clusters. Note. Results from tool by Honorato et al. (2024) [5].

Structural analysis of the docked complex revealed a shape complementarity score (Sc) of 0.72 (calculated using UCSF ChimeraX; range 0–1, with higher values indicating better geometric fit) [30], 18 intermolecular hydrogen bonds (cutoff 3.5 Å), and 6 salt bridges (cutoff 4.0 Å).

### Binding Affinity Prediction

Quantitative affinity prediction using the PRODIGY server identified the top-ranked complex as having an exceptionally strong binding profile. The predicted Gibbs free energy (ΔG) was −21.5 kcal/mol, corresponding to a dissociation constant (Kd) of 1.7 × 10⁻¹⁶ M. Interfacial contact analysis revealed a dense interaction network consisting of 182 total contacts, including 57 hydrophobic–hydrophobic and 55 polar–apolar interactions. The scrambled control peptide yielded a predicted ΔG of −8.3 kcal/mol, substantially weaker than the designed peptide and consistent with nonspecific binding.

**Figure 8.**
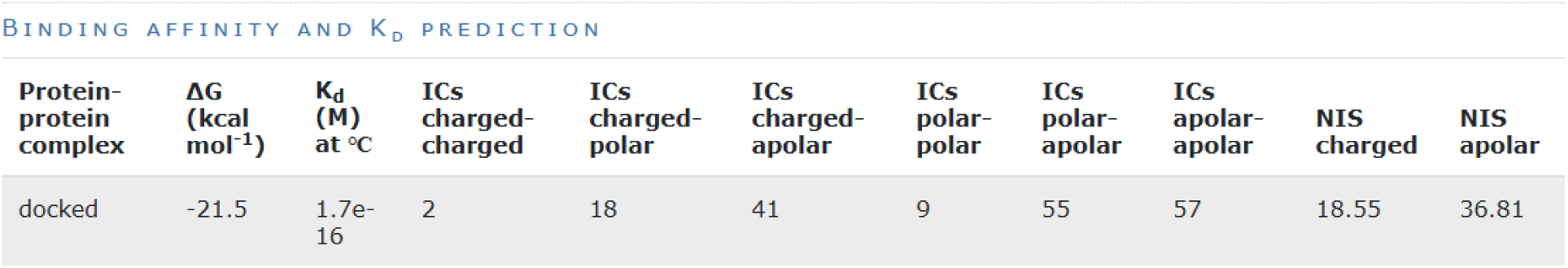
PRODIGY binding affinity and Kd prediction output. Note. Results from tool by Vangone et al. (2016) [16].

**Figure 9.**
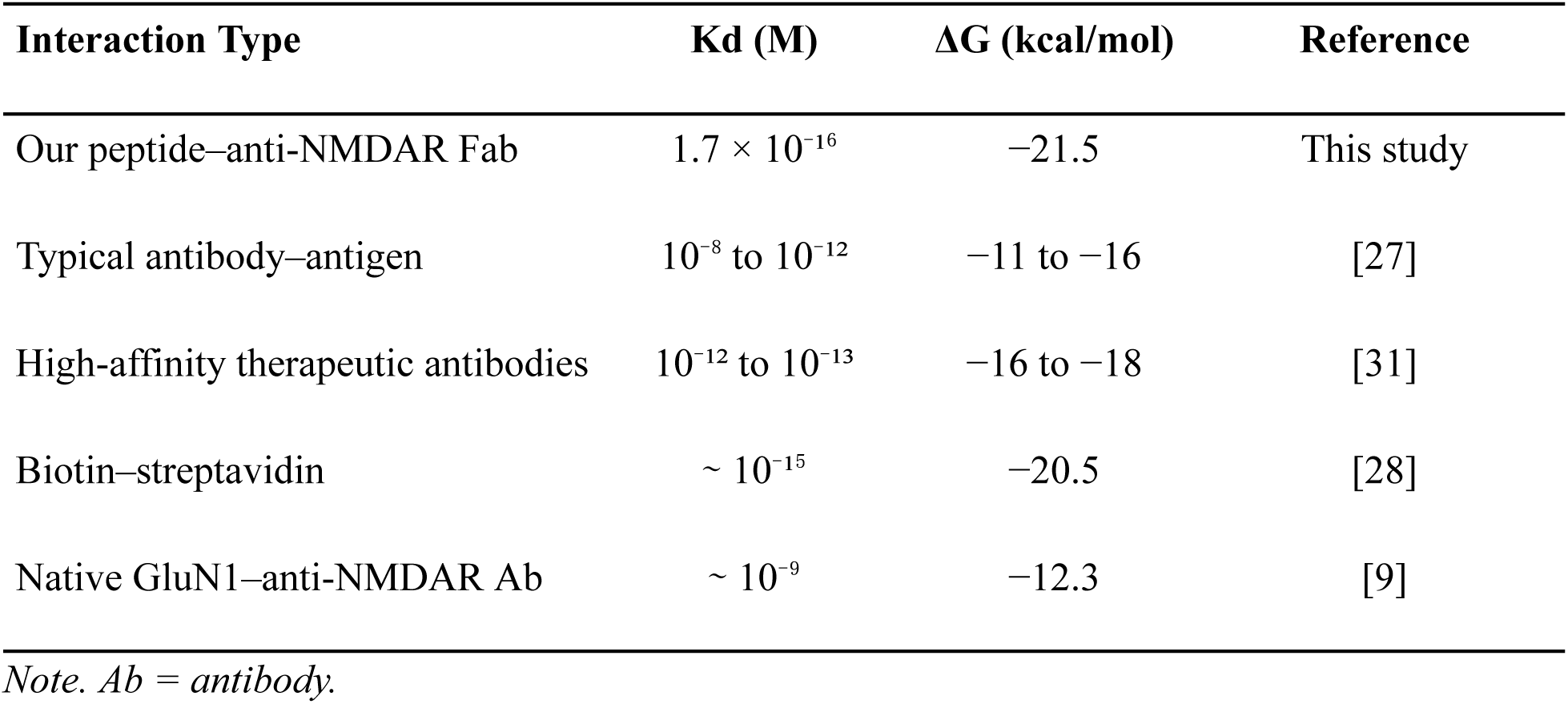
Comparison of predicted binding affinity to literature benchmarks.

The predicted Kd exceeds typical antibody–antigen interactions by several orders of magnitude and approaches the upper limit of known biological binding affinities. As discussed below, PRODIGY predictions may overestimate affinity in peptide–antibody systems, and experimental validation remains essential to confirm these findings.

## Discussion

### Interpretation of Findings

This study presents a fully computational workflow for designing and prioritizing GluN1-mimetic peptide decoys targeting anti-NMDAR antibodies. The top-ranked peptide displayed a favorable predicted binding profile, supported by a large buried surface area (3,255.5 Å²), high shape complementarity (Sc = 0.72), and a dense interfacial contact network (182 contacts). These results suggest that the designed sequence can recapitulate key structural and physicochemical features of the antibody-recognition surface. Binding is predicted to be driven by a combination of extensive hydrophobic packing (57 hydrophobic–hydrophobic contacts) and strong electrostatic complementarity (−310.8 kcal/mol), consistent with the known binding mode of anti-NMDAR antibodies to the GluN1 ATD [8, 9]. The substantial difference in predicted ΔG between the designed peptide (−21.5 kcal/mol) and the scrambled control (−8.3 kcal/mol) supports the specificity of the epitope-mimicry design strategy.

### Comparison with Previous Studies

This work builds on prior computational peptide decoy studies in analogous autoimmune settings. Acetylcholine receptor peptide mimetics in myasthenia gravis have demonstrated preclinical binding activity that correlated with reduced antibody titers in early-phase trials [24]. DNA-binding decoy peptides in systemic lupus erythematosus models showed efficacy in reducing autoantibody-mediated nephritis [18]. In the anti-NMDAR setting specifically, Ehrenreich et al. (2017) identified the GluN1 ATD as the principal antibody target, consistent with our epitope selection [4]. Our computational approach is most comparable to Glasgow et al. (2020), who used engineered ACE2 decoy receptors to potently neutralize SARS-CoV-2 in silico and subsequently validated this strategy experimentally [25]. The current study establishes a similar computational foundation for anti-NMDAR encephalitis while acknowledging that experimental confirmation is needed.

### Limitations

Several important limitations must be acknowledged. First, all results are purely computational and have not been validated experimentally. Predicted binding affinities from PRODIGY should be interpreted as upper-bound estimates rather than true thermodynamic values, as the model was trained on native protein–protein complexes and may not generalize well to designed peptide–antibody systems [32]. The large buried surface area observed (3,255.5 Å², well above the 1,400–2,000 Å² typical of antibody–antigen interfaces [29]) suggests that scoring function bias toward extensive interfaces may inflate the predicted affinity. Second, the HADDOCK docking protocol uses rigid-body approximations during initial orientation generation, which may not fully capture induced-fit effects or the conformational flexibility of short peptides. Third, AlphaFold2 pLDDT scores measure local coordinate confidence rather than thermodynamic stability or aqueous-phase conformational preference; the predicted peptide structure represents a plausible binding-competent state, not necessarily the dominant solution-phase conformation. Fourth, PRODIGY does not fully model entropic penalties associated with peptide folding-upon-binding or the costs of desolvation at highly charged interfaces. Finally, in vivo efficacy would depend on additional factors not modeled here, including blood–brain barrier permeability [34], peptide proteolytic stability, immunogenicity, and pharmacokinetic properties.

### Strengths

Despite these limitations, this study has several notable strengths. The computational pipeline is fully reproducible, utilizing publicly available web servers (HADDOCK 2.4, PRODIGY, ColabFold) and standard software (PyMOL, UCSF ChimeraX). The antibody structure used for docking (PDB: 8ZH7) is a patient-derived humanized monoclonal fragment, making it a clinically relevant binding partner. Epitope selection was grounded in multiple independent lines of crystallographic and mutagenesis evidence, reducing the risk of targeting a non-pathogenic region. The inclusion of a scrambled control peptide enables direct assessment of docking specificity. Physicochemical properties of the lead peptide (net charge +2.09, GRAVY−0.545, predicted good solubility) are favorable for a peptide therapeutic candidate [33]. The pipeline is designed to be modular and scalable, allowing rapid screening of additional variants.

### Future Directions

Several directions are proposed for future work. Experimental validation should begin with surface plasmon resonance (SPR) or isothermal titration calorimetry (ITC) binding assays using recombinant anti-GluN1 Fab fragments to measure actual Kd values. Cell-based internalization assays in cultured hippocampal neurons could assess whether the peptide reduces antibody-induced NMDAR internalization. Peptide stability studies in human serum would characterize proteolytic half-life and identify sites for N-methylation or D-amino acid substitutions to improve proteolytic resistance. Molecular dynamics (MD) simulations would complement the static docking results by capturing peptide flexibility and interface dynamics over time. If binding activity is confirmed in vitro, passive transfer mouse models of anti-NMDAR encephalitis [9] could evaluate in vivo neutralization efficacy. Finally, the computational pipeline described here is directly applicable to other antibody-mediated encephalitides, including anti-LGI1 [39] and anti-AQP4 neuromyelitis optica [37].

## Conclusion

### Summary of Findings

This study employed a fully computational pipeline to design and evaluate a synthetic GluN1-mimetic peptide as a decoy for pathogenic anti-NMDAR antibodies. The lead 41-residue peptide (YSIMNLQNRKLVQVGIYNGTHVIPWDRKIIWPGGETEWPR) achieved a high-confidence AlphaFold2 structure (average pLDDT = 90%), a statistically significant HADDOCK docking cluster (Z-score = −1.9), and a predicted binding affinity substantially stronger than the scrambled control (ΔG = −21.5 vs. −8.3 kcal/mol). Key binding interactions involved the GluN1 epitope residues 351–390 and antibody CDR loops H1, H2, H3, L1, L2, and L3, consistent with the crystallographically characterized binding mode [8, 9].

### Implications

These findings support the feasibility of an epitope-mimicry decoy strategy for anti-NMDAR encephalitis and demonstrate that structure-guided computational screening can efficiently prioritize peptide candidates for further evaluation. The approach offers a potential path toward targeted antibody neutralization without the broad immunosuppression and associated risks of current standard-of-care treatments. The pipeline is scalable and applicable to other autoimmune encephalitides and antibody-mediated neurological diseases.

### Recommendations

We recommend that the lead peptide be advanced to in vitro binding assays using surface plasmon resonance against recombinant anti-GluN1 Fab fragments as the immediate next step. Concurrently, molecular dynamics simulations should be performed to characterize peptide flexibility and interface stability over time. Modifications to improve proteolytic resistance, such as D-amino acid substitutions at cleavage-prone positions, should be explored computationally before synthesis. Ultimately, cell-based and animal model studies will be necessary to assess neutralization efficacy in a physiologically relevant context.

### Closing Statement

This work presents a robust and reproducible computational framework for rational peptide therapeutic design in anti-NMDAR encephalitis and establishes a scalable model applicable to a broader class of antibody-mediated autoimmune disorders. While experimental validation remains essential, the results provide a strong computational foundation for future preclinical development.

## Acknowledgments

We acknowledge the computational resources provided by Google Colab (ColabFold), the HADDOCK web server (Utrecht University), and the PRODIGY server (Utrecht University). We also acknowledge the support of our peers and the publicly available computational resources that made this research possible.

## Author Contributions

**Ravi Shah:** Conceptualization, Methodology, Formal Analysis, Data Curation, Writing – Original Draft, Writing – Review & Editing, Visualization. **Pragyan Misra:** Methodology, Formal Analysis, Writing – Review & Editing, Visualization. **Neeraj Movva:** Methodology, Writing – Review & Editing, Visualization.

